# Empowering Virus Sequences Research through Conceptual Modeling

**DOI:** 10.1101/2020.04.29.067637

**Authors:** Anna Bernasconi, Arif Canakoglu, Pietro Pinoli, Stefano Ceri

**Author notes:** { }.

## Abstract

The pandemic outbreak of the coronavirus disease has attracted attention towards the genetic mechanisms of viruses. We hereby present the Viral Conceptual Model (VCM), centered on the virus sequence and described from four perspectives: biological (virus type and hosts/sample), analytical (annotations and variants), organizational (sequencing project) and technical (experimental technology).

VCM is inspired by GCM, our previously developed Genomic Conceptual Model, but it introduces many novel concepts, as viral sequences significantly differ from human genomes. When applied to SARS-CoV2 virus, complex conceptual queries upon VCM are able to replicate the search results of recent articles, hence demonstrating huge potential in supporting virology research.

In addition to VCM, we also illustrate the data dictionary for patient’s phenotype used by the COVID-19 Host Genetic Initiative. Our effort is part of a broad vision: availability of conceptual models for both human genomics and viruses will provide important opportunities for research, especially if interconnected by the same human being, playing the role of virus host as well as provider of genomic and phenotype information.

## 1. Introduction

Despite the advances in drug and vaccine research, diseases caused by viral infection pose serious threats to public health, both as emerging epidemics (e.g., Zika virus, Middle East Respiratory Syndrome Coronavirus, Measles virus, or Ebola virus) and as globally well-established epidemics (such as Human Immunodeficiency Virus, Dengue virus, Hepatitis C virus). The pandemic outbreak of the coronavirus disease COVID-19, caused by the “Severe acute respiratory syndrome coronavirus 2” virus species SARS-CoV2 (according to the GenBank [35] acronym^1^), has brought unprecedented attention towards the genetics mechanisms of coronaviruses.

Thus, understanding viruses from a conceptual modeling perspective is very important. The sequence of the virus is the central information, along with its annotated parts (known genes, coding and untranslated regions…) and the nucleotides’ variants with respect to the reference sequence for the specific species. Each sequence is identified by a *strain name*, which belongs to a specific virus species. Viruses have complex taxonomies, as they belong to genus, sub-families, and finally families (e.g., Coronaviridae). Other important aspects include the host organisms and isolation sources from which viral materials are extracted, the sequencing project, the scientific and medical publications related to the discovery of sequences; virus strains may be searched and compared intra- and cross-species. Luckily, all these data are made available publicly by various resources, from which they can be downloaded and re-distributed.

Our recent work is focused on data-driven genomic computing, providing contributions in the area of modeling, integration, search and query answering. We have previously proposed a conceptual model focused on human genomics [6], which was based on a central entity Item, representing files of genomic regions. The simple schema evolved into a knowledge graph [5], including ontological representation of many relevant attributes (e.g., diseases, cell lines, tissue types…). The approach was validated through the practical implementation of the integration pipeline META-BASE^2^, which feeds an integrated database, searchable through the GenoSurf^3^ interface [8].

Very recently we have also been involved in the COVID-19 Host Genetics Initiative,^4^ a collaborative effort that aims at joining forces of the broader human genetics community to generate, share, and analyze data to learn the genetic characteristics and outcomes of COVID-19. In this project, we built a conceptual data definition (and related questionnaire) for describing the phenotype of COVID-19, to be used by clinicians who contribute to the project. Thus, we created a conceptually solid definition of the clinical information of patients affected by COVID-19, acting as hosts to the SARS-CoV2 virus. Based on these considerations, in this paper we contribute as follows:

–We propose a new **Viral Conceptual Model (VCM)**, a general conceptual model for describing viral sequences, organized along specific dimensions that highlight a conceptual schema similar to GCM [6];
–Focusing on SARS-CoV2, we show how VCM can be profitably linked to a **phenotype database** with information on COVID-19 infected patients;
–We provide a list of **interesting queries** replicating newly released literature on infectious diseases; these can be easily performed on VCM.

The manuscript is organized as follows: Section 2 overviews current technologies available for virus sequence data management. Section 3 proposes our VCM, while Section 4 shows its possible intersection with a general clinical database. We show examples of applications in Section 5 and review related works in Section 6. Section 7 discloses our vision for future developments.

## 2 Current scenario

The landscape of relevant resources and initiatives dedicated to data collection, retrieval and analysis of virus sequences is shown in Fig. 1. We partitioned the space of contributors by considering: institutions that host data sequences, main sequence databases, tools provided for querying and searching them, and then organizations and tools hosting data analysis interfaces that also connect to viral sequence databases.

**Fig. 1.**
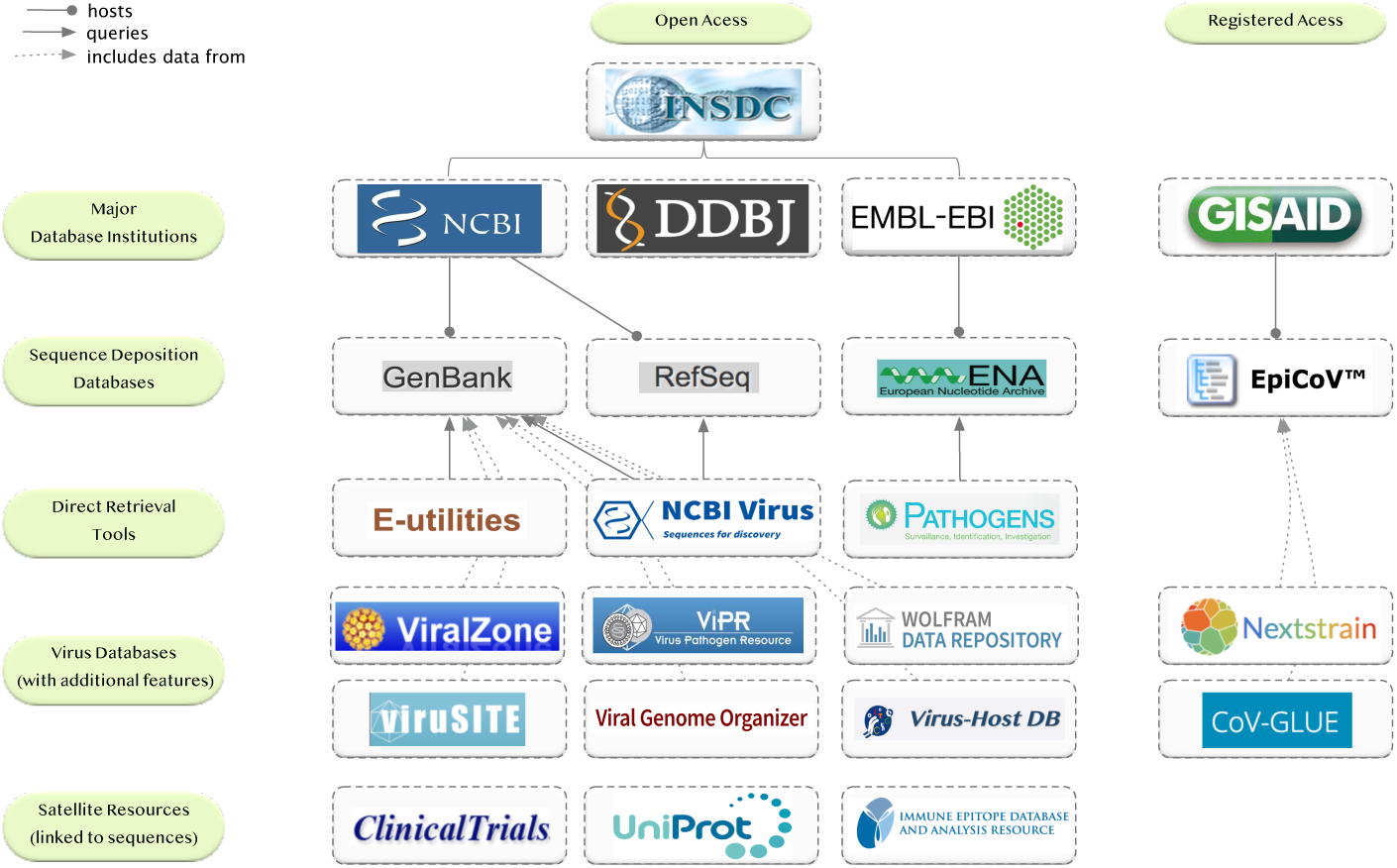
Current relevant resources and initiatives dedicated to data collection, retrieval and analysis of virus sequences, divided by open and registered access.

The three main organizations providing open-source viral sequences are NCBI (US), DDBJ (Japan), and EMBL-EBI (Europe); they operate within the broader contexts provided by the International Nucleotide Sequence Database Collaboration.^5^ NCBI hosts the two, so far, most relevant viral sequence databases: GenBank [35] contains the annotated collection of all publicly available DNA and RNA sequences; RefSeq [28] provides a stable reference for genome annotation, gene identification and characterization, and mutation/polymorphism analysis. GenBank is continuously updated thanks to the abundant sharing of multiple laboratories and data contributors around the world (note that SARS-CoV2 nucleotide sequences have increased from about 300 around the end of March 2020, to 1,624 as of April 27th).

EMBL-EBI hosts the European Nucleotide Archive [1], which has a broader scope, accepting submissions of nucleotide sequencing information, including raw sequencing data, sequence assembly information and functional annotations.

Several tools are available for querying and searching these databases; among them, E-utilities [34], NCBI Virus^6^ [19], and Pathogens^7^ are tools and portals directly provided by the INSDC institutions for supporting the access to their viral resources, however lacking possibility of querying based on annotations and variants.

A number of databases and data analysis tools refer to these viral sequences databases. We mention: ViralZone [20] by the SIB Swiss Institute of Bioinformatics, which provides access to SARS-CoV2 proteome data as well as cross-links to complementary resources; the Virus Pathogen Database and Analysis Resource (ViPR, [31]), an integrated repository of data and analysis tools for multiple virus families, supported by the Bioinformatics Resource Centers program; viruSITE [39], an integrated database for viral genomics; the Viral Genome Organizer,^8^ implemented by the Canadian Viral Bioinformatics Research Centre, focusing on search for sub-sequences within genomes.

While the INSDC consortium provides full open access to sequences, the GISAID Initiative [13, 37] was created in 2008 with the explicit purpose of offering an alternative to traditional public-domain data archives, as many scientists hesitated to share influenza data due to their legitimate concern about not being properly acknowledged, among others. GISAID hosts EpiFlu™, a large sequence database, which started its mission for influenza data and is now expanding with EpiCoV™ having a particular focus on the SARS-CoV2 pandemic (12,645 sequences for SARS-CoV2 on April 27th). Some interesting portals have become interfaces to GISAID data with particular focuses: NextStrain [18] overviews emergent viral outbreaks based on the visualization of sequence data integrated with geographic information, serology, and host species; CoV-GLUE,^9^ part of the GLUE suite [38], contains a database of replacements, insertions and deletions observed in sequences sampled from the pandemic.

Many other resources link to viral sequence data, including: drug databases, particularly interesting as they provide information about clinical studies (see ClinicalTrials^10^), protein sequences databases (e.g., UniProtKB/Swiss-Prot [32]), and cell lines databases (e.g., Cellosaurus [3]).

## 3 Conceptual modeling for viral genomics

We previously proposed the Genomic Conceptual Model (GCM, [6]), an Entity-Relationship diagram that recognizes a common organization for a limited set of concepts supported by most genomic data sources, although with different names and formats. The model is centered on the Item entity, representing an elementary experimental file of genomic regions and their attributes. Four views depart from the central entity, recalling a classic star-schema organization that is typical of data warehouses [7]; they respectively describe: i) the *biological* elements involved in the experiment: the sequenced sample and its preparation, the donor or patient; ii) the *technology* used in the experiment, including a specific assay (i.e., technique); iii) the *management* aspects: the projects/organizations involved in the preparation and production; iv) the *extraction* parameters used for internal selection and organization of items.

GCM is employed as a driver of integration pipelines for genomic datasets, fuelling user search-interfaces such as GenoSurf [8]. Lessons learnt from that experience include the benefits of having: a central *fact* entity that helps structuring the search; a number of surrounding *dimensions* capturing organization, biological and experimental conditions to describe the facts; a data layout that is easy to learn for first-time users and that helps the answering of practical questions (as demonstrated in [4]).

We hereby propose the Viral Conceptual Model (VCM), which is influenced by our past experience with human genomes. There are significant differences between the two conceptual models. The human DNA sequence is long (3 billions of base pairs) and has been understood in terms of *reference genomes* (named h19 and GRCh38) to which all other information is referred, including genetic and epigenetic signals. Instead, viruses are many, their sequences are short (order of thousands of base pairs) and each virus has its own reference sequence; moreover, virus sequences are associated to a host sample of another species.

With a bird eye’s view, the VCM conceptual model is centered on the Sequence entity that describes individual virus sequences; sequences are analyzed from the *biological* perspective (Hostsample and Virus), the *tech- nological* perspective (ExperimentType), and the *organizational* perspective (SequencingProject). Two other entities, Annotation and Variant, represent an *analytical* perspective of the sequence, allowing to analyze its characteristics, its sub-parts, and the differences with respect to reference sequences for the specific virus species. We next illustrate the central entity and the four perspectives.

### Central entity

A viral Sequence can regard either DNA or RNA; in either cases, databases and sequencing data write the sequence as a DNA *NucleotideSequence* (i.e., guanine (G), adenine (A), cytosine (C), and thymine (T)^11^) that has a specific *Strand* (positive or negative), *Length* (typically thousands), and a percentage of read G and C bases (*GC%*). Each sequence is uniquely identified by an *AccessionID*, which is retrieved directly from the source database (GenBank’s are usually formed by two capital letters, followed by six digits, GISAID by the string “EPI ISL” and six digits). Sequences can be complete or partial (as encoded by the boolean flag *IsComplete*) and they can be a reference sequence (stored in RefSeq) or a regular one (encoded by *IsReference*). In the latter case, sequences have a corresponding *StrainName* assigned by the sequencing lab, somehow hard-coding relevant information (e.g., hCoV-19/Nepal/61/2020 or 2019-nCoV PH nCOV 20 026).

### Technological perspective

The sequence derives from one experiment or assay, described in the ExperimentType entity (cardinality is 1:N from the dimension towards the fact). It is performed on biological material analysed with a given *SequencingTechnology* platform (e.g., Illumina Miseq) and an *AssemblyMethod*, collecting algorithms that have been applied to obtain the final sequence, for example: BWA-MEM, to align sequence reads against a large reference genome; BCFtools, to manipulate variant calls; Megahit, to assemble NGS reads. Another technical measure is captured by *Coverage* (e.g., 100X or 77000x).

### Biological perspective

Each sequence belongs to a specific Virus, which is described by a complex taxonomy. The most precise definition is the *Species-Name* (e.g., Severe acute respiratory syndrome coronavirus 2), corresponding to a *SpeciesTaxonID* (e.g., 2697049), related to a simpler *GenBankAcronym* (e.g., SARS-CoV2) and to many comparable forms, contained in the *EquivalentList* (e.g., 2019-nCoV, COVID-19, SARS-CoV-2, SARS2, Wuhan coronavirus, Wuhan seafood market pneumonia virus, …). The species belongs to a *Genus* (e.g., Betacoronavirus), part of a *SubFamily* (e.g., Orthocoronavirinae), finally falling under the most general category of *Family* (e.g., Coronaviridae). Each virus species corresponds to a specific *MoleculeType* (e.g., genomic RNA, viral cRNA, unassigned DNA), which has either double- or single-stranded structure; in the second case the strand may be either positive or negative. These possibilities are encoded within the *IsSingleStranded* and *IsPositiveStranded* boolean variables. An assay is performed on a tissue extracted from an organism that has hosted the virus for an amount of time; this information is collected in the HostSample entity. The host is defined by a *Species*, corresponding to a *SpeciesTaxonID*, usually represented using the NCBI Taxonomy [14] (e.g., 9606 for Homo Sapiens). The sample is extracted on a *CollectionDate*, from an *IsolationSource* that is a specific host tissue (e.g., nasopharyngeal or oropharyngeal swab, lung), in a certain location identified by the quadruple *OriginatingLab* (when available), *Region, Country*, and *GeoGroup* (such as continent). Both entities of this perspective are in 1:N cardinality with the Sequence.

### Organizational perspective

The entity SequencingProject describes the management aspects of the production of the sequence. Each sequence is connected to a number of studies, usually represented by a research publication (with *Authors, Title, Journal, PublicationDate* and eventually a *PubMedID* referring to the most important biomedical literature portal^12^). When a study is not available, just the *SequencingLab* and *SubmissionDate* are provided. In rare occasions, a project is associated with a *PopSet* number, which identifies a collection of related sequences derived from population studies (submitted to GenBank), or with a *BioProjectID* (an identifier to the BioProject external database^13^). We also include the name of *DatabaseSource*, denoting the organization that primarily stores the sequence. In this perspective all cardinalities are 1:N as sequences can be part of multiple projects; conversely, sequencing projects contain various sequences.

### Analytical perspective

This perspective allows to store information that are useful during the secondary analysis of genomic sequences. Annotations include a number of sub-sequences representing segments (defined by *Start* and *Stop* coordinates) of the original sequence with a particular *FeatureType* (e.g., gene, peptide, coding DNA region, or untranslated region, molecule patterns such as stem loops and so on), the recognized *GeneName* to which it belongs (e.g., gene “E”), the *Product* it concurs to produce (e.g., leader protein, nsp2 protein, RNA-dependent RNA polymerase, membrane glycoprotein, envelope protein…), and eventually an *ExternalReference* when the protein is present in a separate database such as UniProtKB. The Variant entity contains sub-sequences of the main Sequence that differ from the reference sequence of the same virus species. They can be identified with respect to the reference one, just by using the *AltSequence* (i.e., the nucleotides used in the analyzed sequence at position *Start* coordinate for an arbitrary *Length*, typically just equal to 1) and a specific *Type*, which can correspond to insertion (INS), deletion (DEL), Single-nucleotide polymorphism (SNP) or others. The content of the attributes of this entity is not retrieved from existing databases; instead it is computed in-house by our procedures. Indeed, we use the well known dynamic programming algorithm of Needleman-Wunsch [26], that computes the optimal alignment between two sequences. From a technical point of view, we compute the pair-wise alignment of every sequence to the reference sequence of RefSeq (NC 045512); from such alignment we then extract all insertions, deletions, and substitutions that transform (edit) the reference sequence into the considered sequence. A similar computation is performed within CoV-GLUE (http://cov-glue.cvr.gla.ac.uk/).

## 4 Phenotype Data Dictionary for COVID-19 Host Genetic Initiative

After the spread of COVID-19 pandemia, several informal consortia have been created to foster international cooperation among researchers. We participate to the COVID-19 Host Genetics Initiative,^14^ aiming at *bringing together the human genetics community to generate, share and analyze data to learn the genetic determinants of COVID-19 susceptibility, severity and outcomes*. In this setting, we are coordinating the production of a data dictionary for the phenotype definition, which will be used as a reference by participating institutions, hosted by EGA [15], the European Genome-phenome Archive of EMBL-EBI.^15^

The dictionary, illustrated in Fig. 3, contains patient phenotype information, collected at admission and during the course of hospitalizations (hosted by a given Hospital); each patient can be connected to a virus sequence (in that case, she is the host organism providing the HostSample of VCS) and can have multiple Encounters. For ease of visualization, attributes are clustered within *Attribute Groups*, indicated with white squares instead of black circles. Note that the dictionary representation deviates from a classic Entity-Relationship diagram as some attribute groups would typically deserve the role of entity; however, this simple format allows an easy mapping of the dictionary to questionnaires and an implementation by EGA in the form of spreadsheet.

**Fig. 2.**
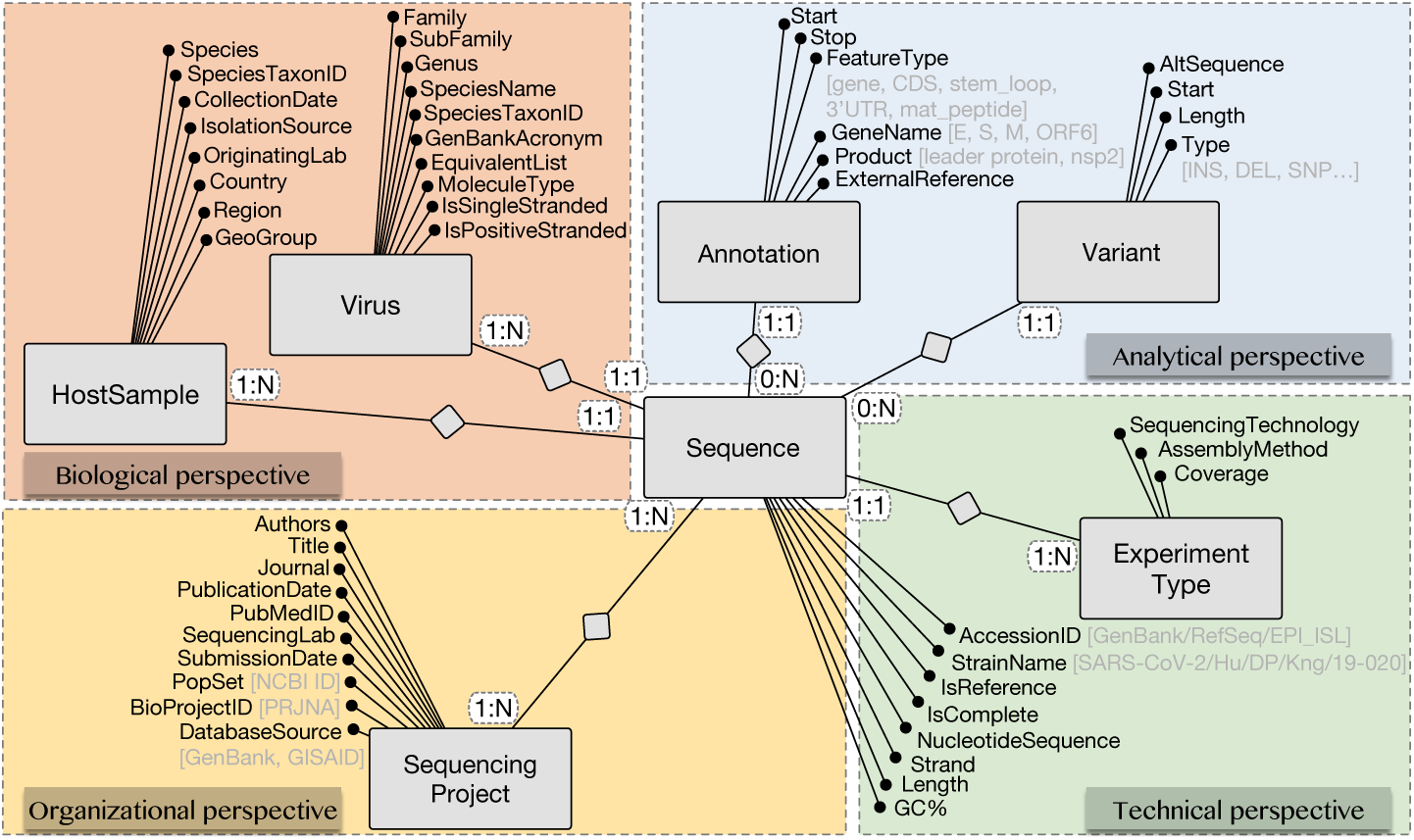
The Viral Conceptual Model: the central fact Sequence is described by four different perspectives (biological, technical, organizational and analytical).

**Fig. 3.**
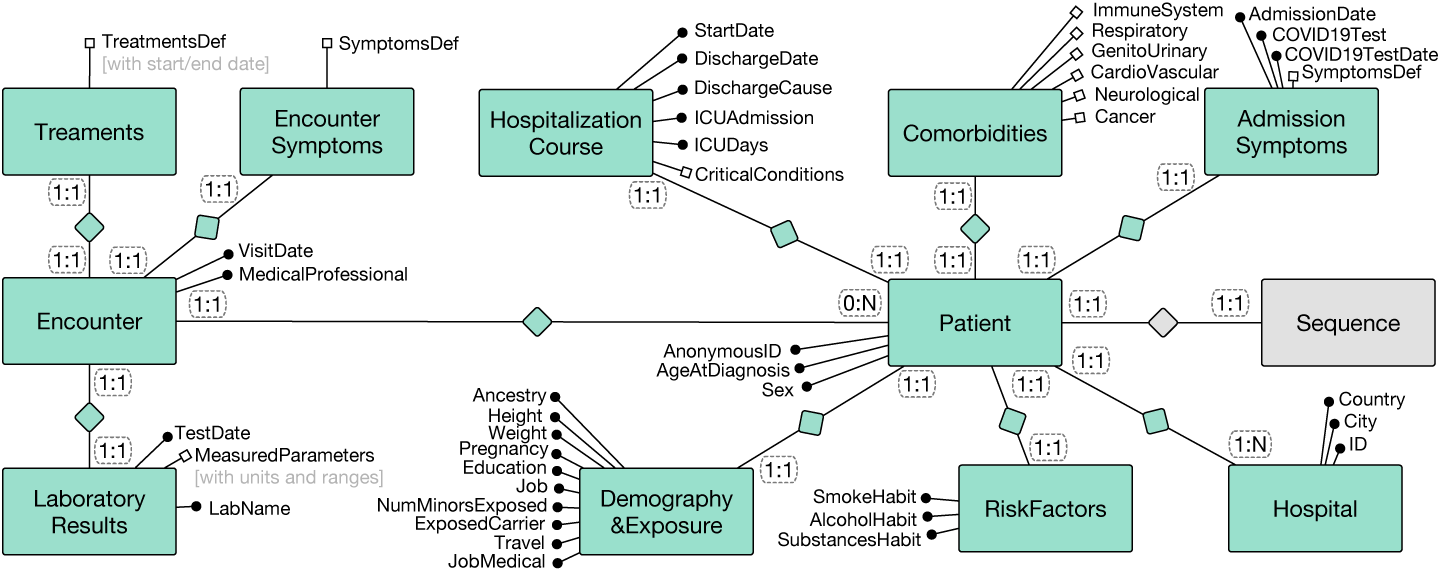
COVID-19 Phenotype Data Dictionary.

Attribute groups of patients describe: Demography&Exposure, Risk-Factors, Comorbidities, AdmissionSymptoms, HospitalizationCourse; attribute groups of encounters describe: EncounterSymptoms, Treatments, LaboratoryResults. Attributes within groups can be further clustered within subgroups; for instance, Comorbidities include the subgroups *ImmuneSystem, Respiratory, GenitoUrinary, CardioVascular, Neurological, Cancer*. The data dictionary includes two possible uses in further analysis (i.e., the course of hospitalization and longitudinal studies); for these uses we set each attribute to either mandatory or optional.

## 5 Answering complex biological queries

In addition to very general questions that can be easily asked through our conceptual model (e.g., retrieve all viruses with given characteristics), in the following we propose a list of interesting application studies that could be backed by the use of our conceptual model. In particular, they refer to SARS-CoV2 virus as it is receiving most of the attention of the scientific community. Fig. 4 represents the reference sequence of SARS-CoV2,^16^ highlighting the major structural sub-sequences that are relevant for the encoding of proteins and other functions. It has 56 region Annotations, of which Fig. 4 represents only the 11 genes (ORF1ab, S, ORF3a, E, M, ORF6, ORF7a, ORF7b, ORF8, N, ORF10) plus the RNA-dependent RNA polymerase enzyme, with approximate indication of the corresponding coordinates. We next describe biological queries supported by VCM, from the easiest to the most complex ones, typically suggested by existing studies.

**Fig. 4.**
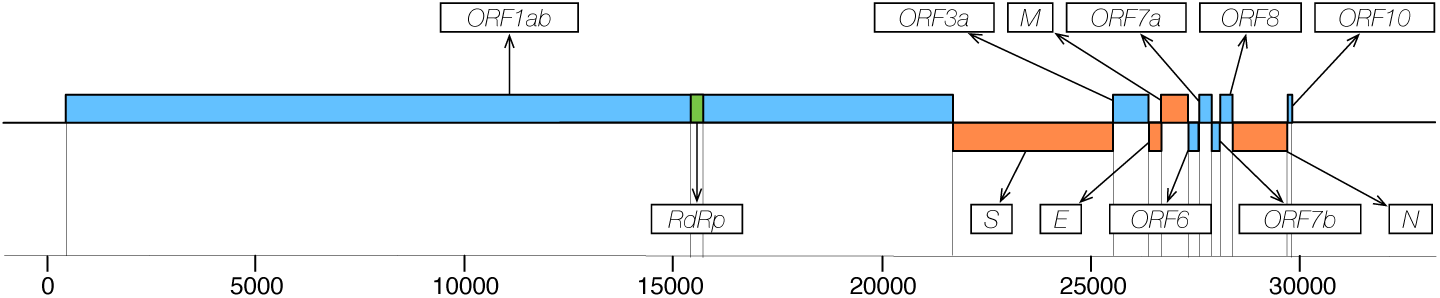
Location of major structural protein-encoding genes (redboxes; S = Spike glycoprotein, E = Envelope protein, M = Membrane glycoprotein, N = Nucleocapsid phospoprotein), accessory protein ORFs = Open Reading Frames (blueboxes), and RNA-dependent RNA polymerase (RdRp) on the sequence of the SARS-CoV2.

**Q1**. The most common variants found in SARS-CoV2 sequences can be selected for US patients; the query can be performed only on specific genes.

**Q2**. COVID-19 European patients affected by a SARS-CoV2 virus can be selected when they have a specific variant on the first gene (ORF1ab), indicated by using the triple *<*start, reference allele, alternative allele*>*. Patients can be distributed according to their country of origin. This conceptual query is illustrated in Fig. 5, where selected attribute values are specified in red, in place of attribute names in the ER model; values in Variant are one possible example. *Country* is in blue as samples will be distributed according to such field.

**Fig. 5.**
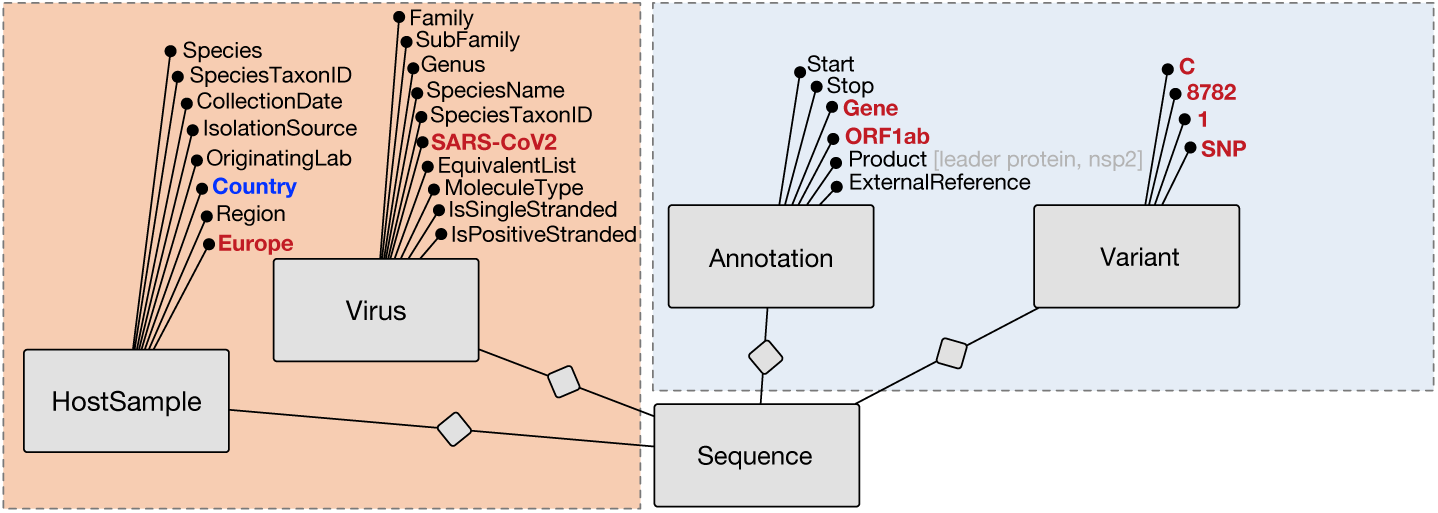
Visual representation of query **Q2**.

**Q3**. According to [9], E and RdRp genes are highly mutated and thus crucial in diagnosing COVID-19 disease; first-line screening tools of 2019-nCoV should perform an E gene assay, followed by confirmatory testing with the RdRp gene assay. Conceptual queries are concerned with retrieving all sequences with mutations within genes E or RdRp and relating them to given hosts, e.g. humans affected in China.

**Q4**. Tang *et al*. [41] claim that there are two clearly definable “major types” (S and L) of SARS-CoV2 in this outbreak, that can be differentiated by transmission rates. Intriguingly, the S and L types can be clearly distinguished by just two tightly linked SNPs at positions 8,782 (within the ORF1ab gene from C to T) and 28,144 (within ORF8 from T to C). Then, queries can correlate these SNPs to other variants or the outbreak of COVID-19 in specific countries (e.g., [16]).

**Q5**. To inform SARS-CoV2 vaccine design efforts, it may be needed to track antigenic diversity. Typically, pathogen genetic diversity is categorised into distinct *clades* (i.e., a monophyletic group on a phylogenetic tree). These clades may refer to ‘subtypes’, ‘genotypes’, or ‘groups’, depending on the taxonomic level under investigation. In [16], specific sequence variants are used to define clades/haplogroups (e.g., the *A group* is characterized by the 20,229 and 13,064 nucleotides, originally C mutated to T, by the 18,483 nucleotide T mutated to C, and by the 8,017, from A to G). VCM supports all the information required to replicate the definition of SARS-CoV2 clades requested in the study. Fig. 6 illustrates the conjunctive selection of sequences with all four variants corresponding to the *A clade group* defined in [16] and the resulting retrieved sequences.

**Fig. 6.**
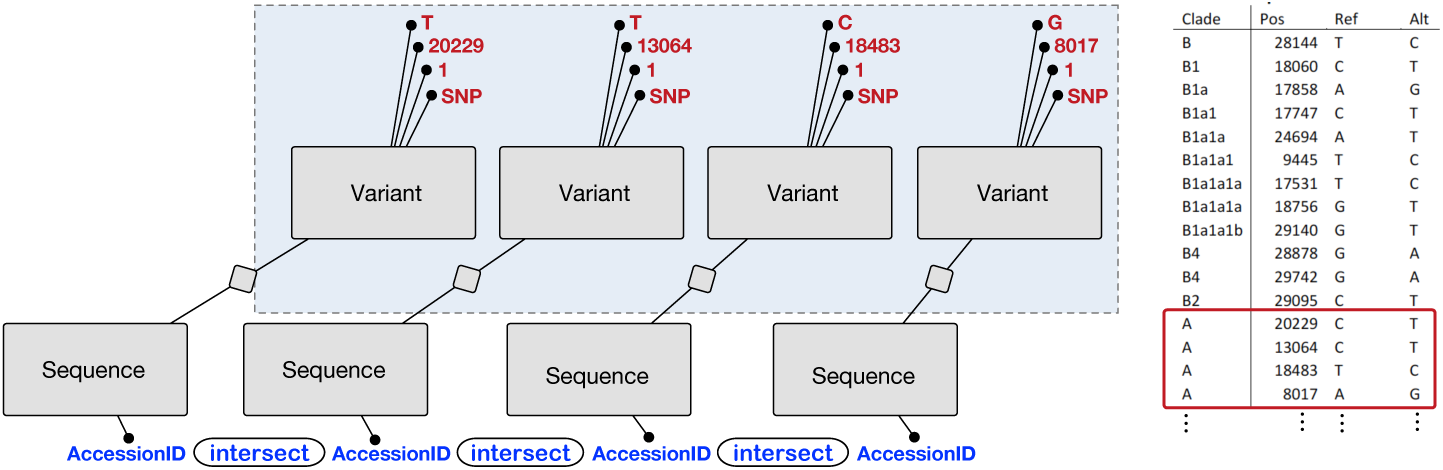
Illustration of the selection predicate for the *A clade group*, used in query **Q5**.

**Q6**. Morais Junior at al. [21] propose a subdivision of the global SARS-CoV2 population into sixteen subtypes, defined using “widely shared polymorphisms” identified in nonstructural (nsp3, nsp4, nsp6, 27 nsp12, nsp13 and nsp14) cistrons, structural (spike and nucleocapsid), and accessory (ORF8) genes. VCM supports all the information required to replicate the definition of such subtypes.

The above examples of complex queries refer to virus sequences and can be answered by VCM (Fig. 2). Due to the pressing interest on SARS-CoV2, we are currently doing an effort to collect SARS-CoV2 sequences and provide a search interface for a first release of a VCM-based query engine.

Even more interesting queries will be enabled by combining phenotypes with virus sequences; along this direction, we also contributed to the data dictionary effort (Fig. 3). When both datasets will be accessible, other more powerful studies will be possible. Some early findings have been already published connecting virus sequences with phenotypes, so far with very small datasets (e.g., [22] with only 5 patients, [24] with 9 patients, and [41] with 103 sequenced SARS-CoV2 genomes). As reaffirmed by these works, there is need for additional comprehensive studies linking the viral sequences of SARS-CoV2 to the phenotype of patients affected by COVID-19. We are confident that in the near future there will be many more studies like [22, 24, 41].

## 6 Related work

The use of conceptual modeling to describe genomics databases dates back to the late nineties, including a functional model for DNA databases named “associative information structure”[27]; a model representing genomic sequences [25]; and a set of data models for describing transcription/translation processes [30]. Later on, a stream of works on conceptual modeling-based data warehouses includes the GEDAW UML Conceptual schema [17], driving the construction of a gene-centric data warehouse for microarray expression measurements; the Genomics Unified Schema [2]; the Genome Information Management System [10], a genome-centric data warehouse; and the GeneMapper Warehouse [12], integrating expression data from a number of genomic sources.

More recently, there has been a solid stream of works dedicated to data quality-oriented conceptual modeling: [33] presents the Human Genome Conceptual Model and [29] applies it to uncover relevant information hidden in genomics data lakes. Conceptual modeling has been mainly concerned with aspects of the *human* genome, even when more general approaches were adopted; in [6] we presented the Genomic Conceptual Model (GCM), describing the metadata associated with genomic experimental datasets available for humans or other model organisms; GCM was essential for driving the data integration pipeline and building search interfaces [8].

In the variety of types of genomic databases [11], several resources are dedicated to viruses [36]; however, very few works relate to conceptual data modeling. Among them, [40] considers host information and normalized geographical location, and [23] focuses on influenza A viruses. The closest work to us, described in [38], is a flexible software system for querying virus sequences; it includes a basic conceptual model^17^. In comparison, VCM covers more dimensions, that are very useful for supporting research queries on virus sequences.

## 7 Conclusions and future developments

This paper responds to an urgent need, understanding the conceptual properties of SARS-CoV2 so as to facilitate research studies. However, the model applies to any type of virus, and will be at the basis for the development of new instruments. In the past, we first presented the conceptual model for human genomics [6], then we developed the Web-based search system GenoSurf [8]; our ongoing effort is to develop a search system for viral conceptual schemas, inspired by GenoSurf.

While the need for data is pressing, there is also a need of conceptually well-organized information. In our broad vision, the availability of conceptual models for both human genomics and viruses will provide important opportunities for research, amplified to the maximum when human and viral sequences will be interconnected by the same human being, playing the role of host of a given virus sequence as well as provider of genomic and phenotype information.

In the future we will continue our modeling and integration efforts for virus genetics in the context of humans, by interacting with the community of scholars who study viruses. We may add more discovery-oriented entities to the model, that could be of use in a future scenario, e.g., a new pandemic offspring. A user researching on diagnosis could ask, for example, what sequence patterns are unique to the whole or sub-part of the database (i.e., do not appear in viruses within the database). Whereas, a user working on vaccine development could be interested in what are the epitopes (i.e., antigen parts to which antibodies attach) that cover the whole database or a partition of it, for MHC types prevalent in different infected humans. Possibly, other dimensions will be necessary, such as drug resistance information and drug resistance-associated mutations.

## Acknowledgements

This research is funded by the ERC Advanced Grant 693174 GeCo (Data-Driven Genomic Computing), 2016-2021. The authors thank Prof. Limsoon Wong for his precious suggestions and inspiration for future works.

## Footnotes

1 SARS-CoV2 is generally identified by the NCBI taxonomy ID 2697049, https://www.ncbi.nlm.nih.gov/Taxonomy/Browser/wwwtax.cgi?id=2697049.

2 META-BASE is available at https://github.com/DEIB-GECO/Metadata-Manager.

3 http://gmql.eu/genosurf/

4 https://www.covid19hg.org/

5 http://www.insdc.org/

6 https://www.ncbi.nlm.nih.gov/labs/virus

7 https://www.ebi.ac.uk/ena/pathogens/

8 https://4virology.net/virology-ca-tools/vgo/

9 http://cov-glue.cvr.gla.ac.uk/

10 http://clinicaltrials.gov/

11 In RNA sequencing databases uracil (U) is replaced with thymine (T).

12 https://www.ncbi.nlm.nih.gov/pubmed/

13 https://www.ncbi.nlm.nih.gov/bioproject/

14 https://www.covid19hg.org/

15 We coordinated about 50 active participants and produced Freeze-1 of the data dictionary, released on April 16, 2020, available at http://gmql.eu/phenotype/.

16 It represents the positive-sense, single-stranded RNA virus (from 0 to the 29903^*th*^ base) of NC 045512 RefSeq complete sequence (*StrainName* “Wuhan-Hu-1”), collected in China from a “Homo Sapiens” HostSample in December 2019; it has been curated by NCBI staff.

17 http://glue-tools.cvr.gla.ac.uk/images/projectModel.png

